# Computational reconstruction of hierarchical cis-regulatory networks reveals synergistic transcription control and disease-associated rewiring

**DOI:** 10.64898/2026.06.24.734159

**Authors:** Xunuo Zhu, Xuheng Zhou, Yufang Zhang, Guoxin Cai, Wenyi Zhao, Binbin Zhou, Jingqi Zhou, Zefang Tang, Jie Liu, Qiang Zhu, Ji Cao, Bo Yang, Xun Gu, Zhan Zhou

## Abstract

Gene regulation emerges from coordinated interactions among dispersed cis-regulatory elements, yet how these elements integrate into functional regulatory networks and collectively regulate gene transcription remains poorly understood. Here, we present ORIGAMI, a multi-omics, gene-centric deep learning framework that reconstructs functional cis-regulatory networks constrained by transcriptional output. ORIGAMI formulates cis-regulatory modeling as a latent graph inference task, which integrates DNA sequence, epigenomic signals, and three-dimensional chromatin priors to infer denoised regulatory graphs that capture functional interactions rather than structural proximity alone. The inferred regulatory graphs exhibit distinct topological regimes, where hierarchical and modular organization encodes cell-state-specific functional demands and enables synergistic transcriptional control. Furthermore, we show that these regulatory architectures undergo measurable state-dependent rewiring across disease contexts. Finally, ORIGAMI accurately predicts the transcriptional consequences of both cis- and trans-regulatory perturbations and links the rearrangement of regulatory architecture to perturbation response. Together, ORIGAMI advances a network-based view of gene regulation and establishes a foundation for virtual cell modeling of regulatory dynamics.

## Main

Gene regulation is fundamentally an emergent property of distributed molecular interactions across multiple genomic scales, in which dispersed candidate cis-regulatory elements (cCREs) collectively coordinate spatiotemporal expression of genes ^1^. CREs are regions of non-coding DNA which reside hundreds to millions of base pairs upstream or downstream of the genes they regulate ^2–5^, and variations within CREs may lead to various diseases by disrupting the normal expression of target genes ^6–8^. Rather than acting independently, cCREs form structured and hierarchical interaction networks that integrate signals across three-dimensional chromatin folding to regulate gene expression ^9–11^. However, due to the complexity and cell type-specificity of the interactions between CREs, deciphering how local and long-range regulatory grammar are integrated into gene-centric cis-regulatory networks remains a central challenge in regulatory genomics ^12–15^.

Experimentally constructing cis-regulatory network suffers from low resolution and high cost, while computationally reconstructing regulatory architectures requires overcoming significant data sparsity and noise. Chromatin conformation capture technologies such as Hi-C only measure physical proximity between genomic segments rather than direct regulatory activity^16^. Regulatory activity assays such as CRISPR screening^17–20^, STARR-seq^21,22^, and massively parallel reporter assays (MPRA)^23,24^, enable systematic evaluation of CREs activity. However, the vast number of cCREs and their context-specific activity ^3^ make comprehensive assessment across all elements and their combinations in different cell types infeasible. Recent deep learning methods have made substantial progress in inferring cis-regulatory grammar and predicting gene expression from genomic and epigenomic inputs^12,19,25–31^, yet most approaches implicitly infer CRE-gene interactions from attention mechanisms or gradient analysis and overlooking the explicit modeling of hierarchical long-range regulation between CREs ^19,25,28–30^. Several graph-based models have begun to incorporate 3D genome information, but they often inherit noise from chromatin contact data and lack mechanisms to distinguish functional interactions from structural proximity^27,31^, limiting their ability to capture causally relevant regulatory relationships.

Here we present ORIGAMI, multiOmics-based cis-Regulatory Interaction and Gene Activity Modeling with hierarchical Inference, to infer functional cis-regulatory networks by explicitly linking network reconstruction to transcriptional regulation. ORIGAMI also means an ancient art of folding paper into decorative shapes, which just like the intricate cis-regulatory network, folding distant regulatory elements together with promoter. Taking multi-omics sequence profiles with three-dimensional chromatin conformation data of cCREs flanking the target gene as input, ORIGAMI jointly reconstructs cis-regulatory networks for each gene and predicts expression using the reconstructed network. Specifically, ORIGAMI reconstructs hierarchical interactions among all the regulatory elements of a target gene using sparse and noisy chromatin contact data as prior interactions. This is achieved by a graph autoencoder that embeds regulatory elements into a latent space, where cCREs with similar regulatory function and multi-omics features are more closer, thereby eliminated noise from prior interactions. Meanwhile, the expression prediction task guides and constrains model to reconstruct true regulatory interactions, which means only interactions that support accurate expression prediction are retained, ensuring that the learned graph captures causal and functional regulatory structure, rather than merely structural chromatin proximity.

Leveraging data across multiple cell types, we show that the regulatory complexity of networks tightly links to functional specialization of cell identity and undergoes targeted rewiring during tumorigenesis. By performing in silico perturbations of cis- and trans-regulatory variants, we found that ORIGAMI accurately predicts perturbation responses and revealed how non-coding variations drive transcriptional reprogramming through the cis-regulatory network rewiring. In a word, ORIGAMI models the hierarchical organization of gene cis-regulation, from local sequence within individual cCREs to long-range, gene-centric coordination among multiple regulatory elements. This computational framework bridges the gap between static multi-omics measurements and dynamic systems biology, providing a generalizable engine for the predictive modeling of virtual cells.

## Results

### ORIGAMI formulates gene regulation as function-constrained latent graph inference

ORIGAMI formulates the deciphering of gene regulation as a graph inference problem. By integrating genomic and epigenomic profiles of cCREs, together with chromatin interaction information, the framework jointly reconstruct cis-regulatory interaction networks and predict downstream gene expression (Fig. 1). ORIGAMI is trained using cell line-specific active cCREs identified by Encyclopedia of DNA Elements (ENCODE) ^3^ as the fundamental units rather than whole genome sequences. This design substantially reduces sequence redundancy and background noise introduced by inactive genomic regions, while focusing the model capacity on regions with regulatory potential to achieve ultra-long range cis-regulatory modeling ^30,32^.

**Fig. 1.**
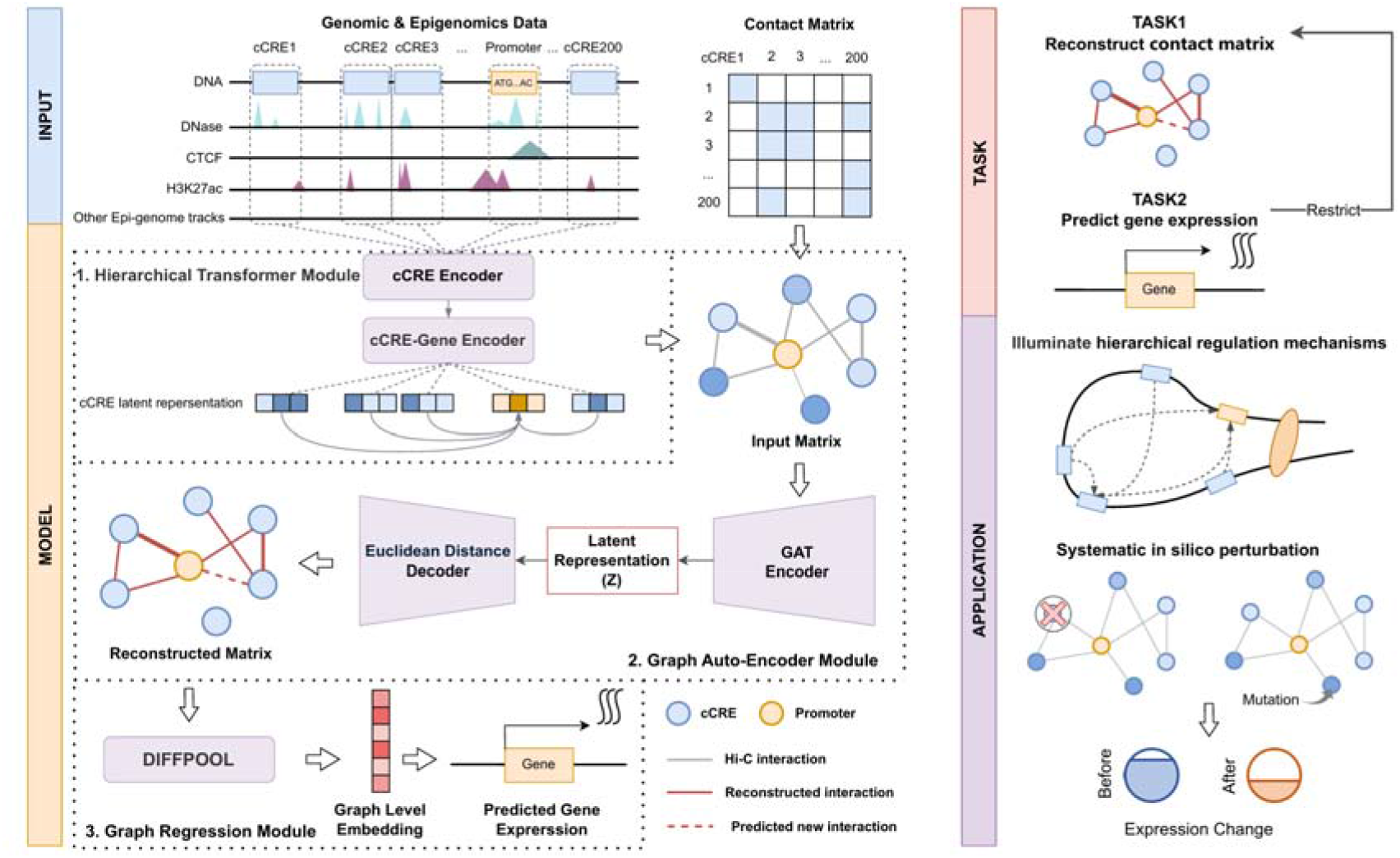
Overview of ORIGAMI. ORIGAMI integrates multi-omics regulatory features and 3D chromatin organization to jointly reconstruct cis-regulatory interaction networks and predict gene expression. (A) For each target gene, base-resolution genome sequence, DNase-seq, and histone/TF ChIP-seq signals are extracted from 200 surrounding cCREs and the promoter, together with a Hi-C-derived contact matrix providing a sparse and noisy prior of regulatory interactions. (B) ORIGAMI consists of three modules. (i) A hierarchical transformer module first encodes representations for each regulatory elements (cCRE encoder) and then aggregates information across all elements (cCRE-gene encoder) to obtain regulatory-aware embeddings. (ii) A graph autoencoder module takes these embeddings and the Hi-C prior as input to reconstruct a denoised cis-regulatory interaction matrix. (3) A graph regression module employs DiffPool to derive a graph-level embedding, which is mapped to target gene expression. (C) The model is trained end-to-end to reconstruct regulatory contact matrices and gene expression, enabling downstream analyses including interpretation of hierarchical regulatory mechanisms and how disruptions in cis/trans-regulation lead to gene dysregulation.

The model is composed of three modules that together capture the multi-scale hierarchical nature of cis-regulatory (Fig. 1B). First, a hierarchical transformer module encodes regulatory information of cCREs and promoter flanking the target gene at two levels: a CRE-centric encoder aggregates base-level genome sequence and epigenomic signals into latent representation for each cCRE, while a regulatory-centric encoder models long-range regulatory relationships of cCREs through genomic position-aware attention mechanism. This design thereby generates the regulatory-aware embeddings that contains both local regulatory grammar and gene-level regulatory coordination.

Second, we initialize the regulatory landscape of each target gene as a graph: regulatory-aware embeddings as node features, and Hi-C-derived chromatin contacts as sparse and noisy prior edges. Then, a graph autoencoder module encodes cCRE latent representations by aggregating 3D genome-neighboring cCRE information and reconstructs the denoised interaction probability of cCRE-cCRE and cCRE-promoter. Importantly, this step transforms noisy and low-resolution chromatin contact data into gene-specific cis-regulatory networks.

Finally, ORIGAMI constrains cis-regulatory network reconstruction task through a graph regression-based gene expression prediction task. The reconstructed network is hierarchically aggregated using differentiable graph pooling (DiffPool), yielding a graph-level gene representation which is subsequently regressed to observed expression values. This coupling of two tasks ensures that only true regulatory interactions, which contribute to accurate expression prediction, are retained, thereby filtering spurious contacts and enhancing biological relevance.

In summary, by explicitly coupling latent graph reconstruction with gene expression prediction, the model learns cis-regulatory networks that encode functional regulatory causality rather than mere chromatin proximity, enabling mechanistic interpretation of non-coding regulatory architecture.

### ORIGAMI infers functional regulatory graphs from sparse chromatin priors

We first asked whether function-constrained graph inference can recover cis-regulatory interactions from noisy chromatin-contact priors in a manner that generalizes across genomic and cellular contexts. To this end, we trained ORIGAMI across nine cell lines whose cell line-specific activity cCRE information and epigenome profiles were available (Supplementary Fig. S1). In cell lines except for K562, chr16 was left out for validation, chr8 and 9 were left out for testing (in-cell type test chromosomes), and all other autosomes were used for training. All autosomes of K562 were left out for testing (cross-cell type test chromosomes) (Supplementary Fig. S2). Especially, neither the genome nor the epigenome information for chr8 and chr9 of K562 had been seen by the model, making them a strictly zero-shot test set.

In the evaluation of the ability to reconstruct cis-regulatory interaction matrices, ORIGAMI achieved consistently high reconstruction accuracy, as quantified by area under the ROC curve (auROC) across held-out chromosomes within the same cell type (in-cell type test chromosomes) (Fig. 2A) and across cell types (cross-cell type test chromosomes) (Fig. 2B). Notably, ORIGAMI maintained robust performance in zero-shot setting, with 0.84 auROC and 0.82 area under the precision-recall curve (auPRC) of K562 chr8 and chr9 (Fig. 2C), suggesting that ORIGAMI can accurately reconstruct cis-regulatory interactions in unseen cell types without additional training. Importantly, ORIGAMI is not limited to recovering physical contacts between genomic fragments, but also captures functionally causal regulatory links. Benchmarking against six baseline methods, ORIGAMI showed superior performance in prioritizing CRISPR-validated K562 cCRE-gene pairs (Fig. 2D). In particular, ORIGAMI’s performance far exceeds the performance achieved using merely Hi-C-derived prior information (cCRE contact matrix), highlighting the necessity of aggregating contextual regulatory signals across multiple cis-regulatory elements.

**Fig. 2.**
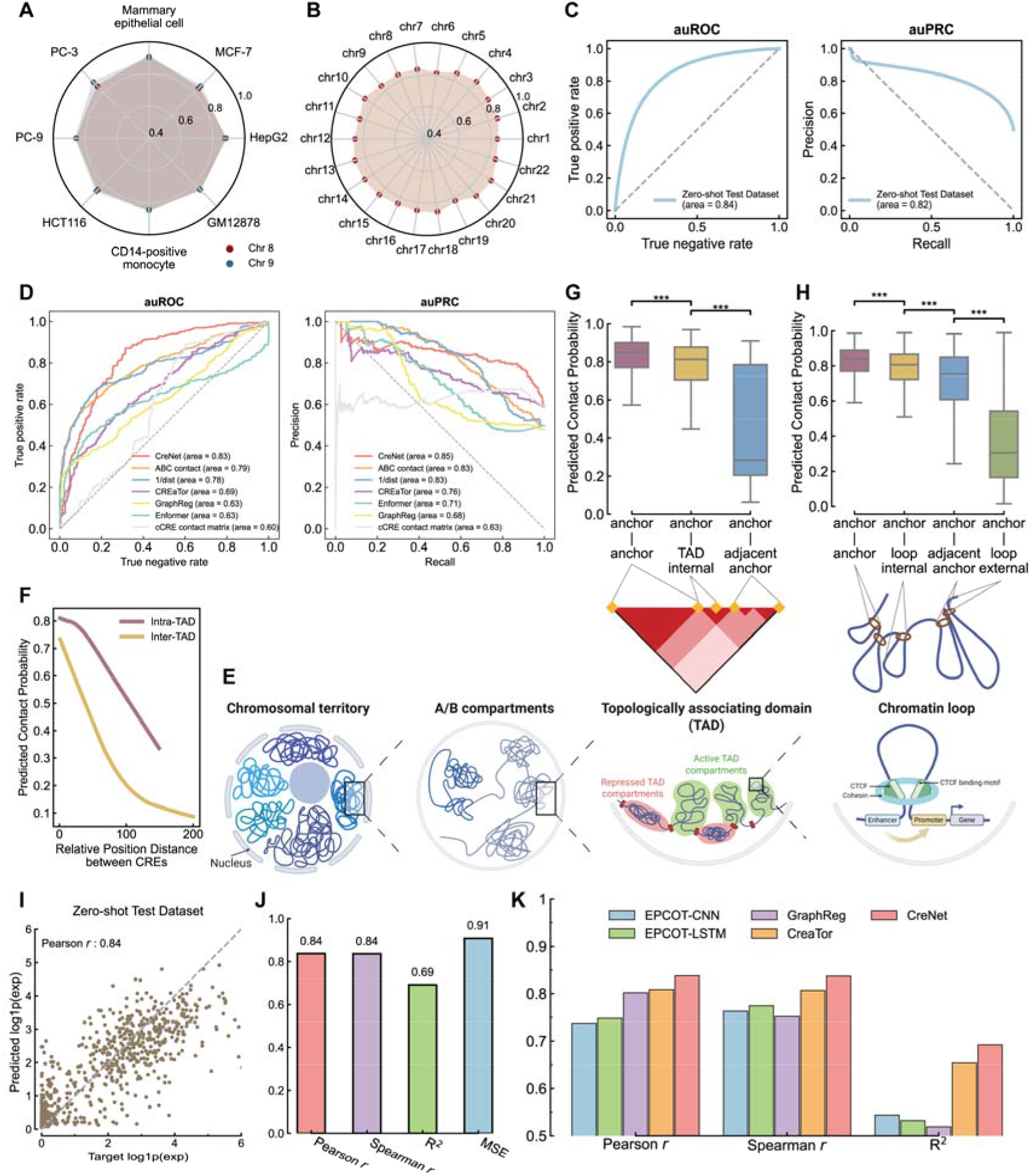
Accurate gene cis-regulatory network reconstruction and expression prediction with ORIGAMI. (A-B) Area under the ROC curve (auROC) evaluating the reconstruction performance of the cis-regulatory interaction networks, assessed on in-cell type test chromosomes (A) and cross-cell type (K562) test chromosomes (B). (C) ROC (left) and precision-recall (PR) curves (right) evaluating the reconstruction performance of the cis-regulatory interaction networks, assessed on zero-shot test chromosomes (K562 chr8 & chr9). (D) Comparison of CRISPR-validated cCRE-gene interaction prioritization across ORIGAMI and six baseline methods, evaluated by auROC (left) and auPRC (right). (E) Schematic illustration of hierarchical 3D genome organization. (F) Predicted contact probability decays with increasing genomic distance, stratified by intra- and inter-TAD interactions. The mean contact probabilities at different distance are linear regressed using locally weighted linear regression to capture distance-dependent interaction decay. (G-H) The reconstructed cis-regulatory network captures hierarchically genome organizations, including TADs (G) and chromatin loops (H). (G) Predicted contact probabilities for interactions between adjacent anchors, within the same TAD, and between distant anchors. (H) Predicted contact probabilities between loop anchors and loop-internal and external regions. (I) Scatter plot showing high consistency between predicted and target gene expression in zero-shot test chromosomes. (J) Pearson *r*, Spearman *r*, R^2^, and mean squared error (MSE) demonstrate expression prediction performance on zero-shot test chromosomes. (K) Benchmark comparison of gene expression prediction performance between ORIGAMI and four baseline methods. *P*-values were calculated using Student’s *t*-test (* *P* < 0.05; ** *P* < 0.01; *** *P* < 0.001).

Mechanistically, ORIGAMI’s performance arises from the joint modeling of regulatory representation, prior graph structure, and functional constraint (Supplementary Fig. S3). Removing either expression guidance or Hi-C priors impaired cCRE-gene pair prioritization, demonstrating that functional constraint and structural initialization contribute complementary information for regulatory graph inference. Meanwhile, removing Hi-C prior caused an even larger degradation of performance, underscoring its role as a structural scaffold. In terms of model architecture, replacing the graph autoencoder with a transformer-only architecture led to the most substantial degradation in cCRE-gene pairs prioritization, demonstrating that explicit latent graph inference is critical for denoising and propagating regulatory signals beyond pairwise attention. Finally, removing the regulatory-centric transformer or replacing DiffPool with mean pool resulted in moderate but consistent declines, highlighting their roles in capturing long-range dependencies and preserving hierarchical structure.

Beyond pairwise accuracy, the reconstructed matrices recapitulated known biophysical constraints of genome folding (Fig. 2E). Topologically associating domains (TADs) are self-interacting genomic regions in which cis-regulatory elements preferentially contact other elements within the same domain, resulting in higher intra-domain contact frequencies than inter-domain interactions ^33^. Consistent with this topological structure, ORIGAMI captured the characteristic decay of contact probability with increasing genomic distance and clear stratification between intra- and inter-TAD interactions (Fig. 2F). We further leveraged previously defined K562 TAD boundaries ^34^ to evaluate whether the model learned domain insulation behavior. As shown in Figure 2G, CRE pairs that locate at the boundary of same TAD (anchor-anchor) were assigned higher contact probability compared to contacts either between boundary and TAD inner region (anchor-TAD internal), or unmatched boundaries spanning adjacent TADs (anchor-adjacent anchor). Similarly, at a more refined level of chromatin loops, ORIGAMI confers significantly higher contact probabilities for interactions within the same loop and between anchors of same loop, far exceeding those for interactions spanning adjacent loops (Figure 2H). These patterns indicate that ORIGAMI implicitly captures multi-scale topological constraints of 3D genome organization.

More crucially, this reliable network reconstruction successfully translates into improved gene expression prediction. ORIGAMI achieved high concordance between predicted and observed expression levels on zero-shot test dataset (Pearson *r* = 0.84) (Fig. 2I-J). Compared with four retrained state-of-the-art baselines, ORIGAMI consistently achieved the best performance across multiple metrics (Fig. 2K). Together, these findings demonstrate that ORIGAMI reconstructs functionally-grounded regulatory hierarchies rather than isolated enhancer-gene pairs, thus bridging noncoding genome organization with transcriptional output.

### Reconstructed regulatory graphs display distinct topological regimes

To quantitatively analyze the systemic organization governing gene regulation, we transformed the continuous interaction matrices inferred by ORIGAMI into cis-regulatory networks, in which nodes correspond to cCREs (labeled as 0-99 and 101-200) and promoter (labeled as ‘P’), and weighted edges represent predicted contact probabilities. For each gene, edges exceeding a defined confidence weight and nodes with shortest paths to the promoter no greater than 3 were retained to form a regulatory network, enabling direct comparison of network topology across genes (Fig. 3A).

**Fig. 3.**
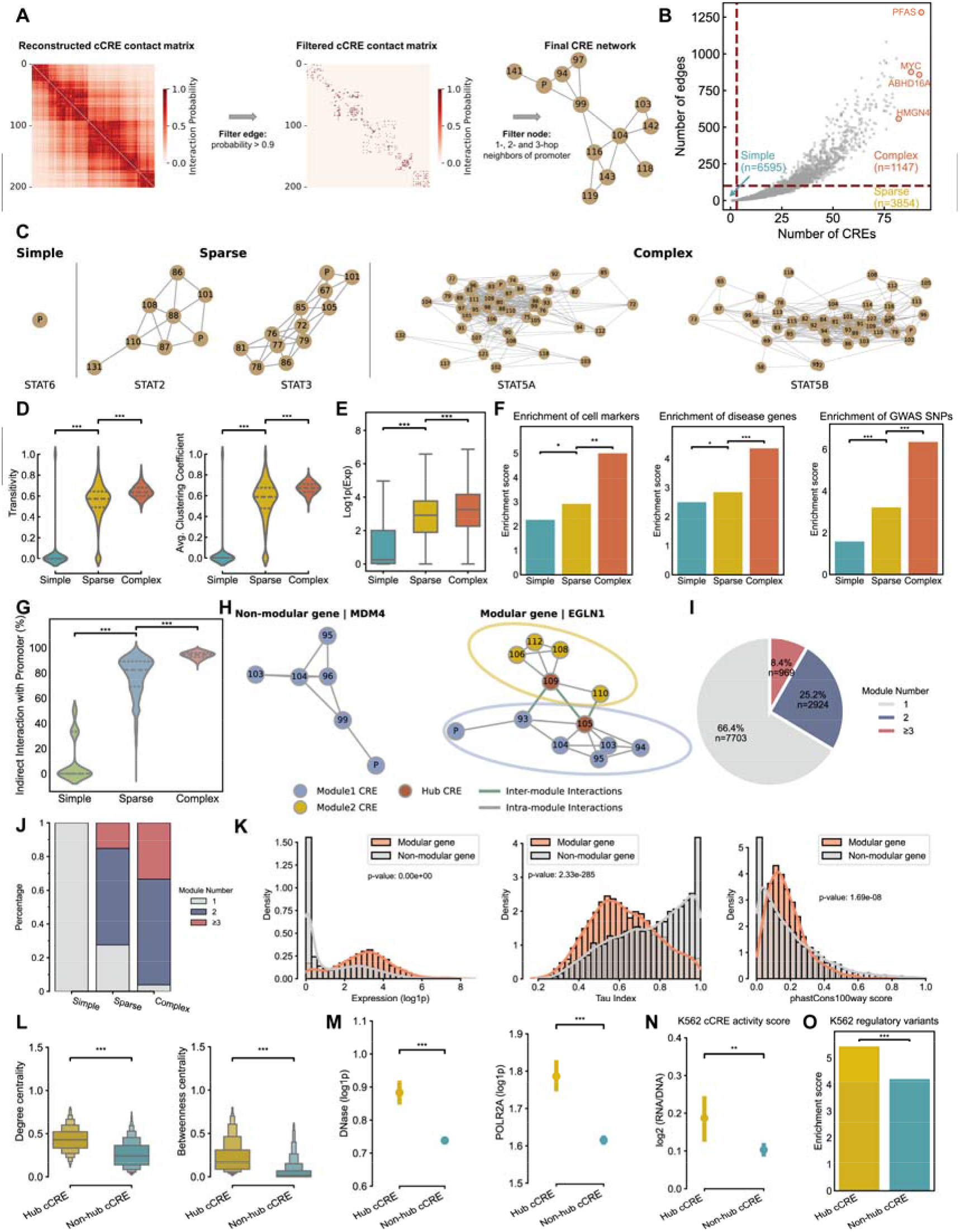
Complex cis-regulatory networks are modularly organized. (A) Workflow for deriving cis-regulatory networks from reconstructed interaction matrices. (B) Classify the K562 cis-regulatory network into simple, sparse, and complex network based on the number of cCREs (nodes) and interactions (edges). Representative genes with complex networks are highlighted. (C) Representative cis-regulatory networks of STAT family members in K562, illustrating simple (STAT6), sparse (STAT2 and STAT3), and complex (STAT5A and STAT5B) regulatory architectures. ‘P’ denotes the promoter, and numbered nodes are CREs. (D) Comparison of topological properties across three types of networks. *P*-values were calculated using Student’s *t*-test (* *P* < 0.05; ** *P* < 0.01; *** *P* < 0.001). (E) Comparison of target gene expression levels across three types of networks. *P*-values were calculated using Student’s *t*-test (* *P* < 0.05; ** *P* < 0.01; *** *P* < 0.001). (F) Left and Middle: Enrichment of cell markers and leukemia-associated disease genes in genes with simple, sparse and complex networks. Right: Enrichment of GWAS SNPs in CREs within simple, sparse and complex networks. *P*-values were calculated using binomial test (* *P* < 0.05; ** *P* < 0.01; *** *P* < 0.001). (G) Fraction of indirect CRE-promoter interactions across three types of networks. *P*-values were calculated using Student’s *t*-test (* *P* < 0.05; ** *P* < 0.01; *** *P* < 0.001). (H) Examples illustrating cis-regulatory networks with non-modular and modular structure. In modular network, green line represents inter-module interaction and gray line represents intra-module interaction. The hub CREs were further identified within the modular organized networks, labeled as red nodes. (I) Pie plot showing the percentage of cis-regulatory networks stratified by the module numbers. (J) Distribution of module numbers within simple, sparse, and complex networks. (K) Comparison between modular and non-modular genes in terms of expression level (left), Tau tissue-specificity index (middle), and phastCons100way conservation score (right). *P*-values were calculated using Student’s *t*-test (* *P* < 0.05; ** *P* < 0.01; *** *P* < 0.001). (L-N) Comparison between hub and non-hub cCREs in modular genes in terms of centrality measures including degree centrality and betweenness centrality (L), epigenomic activity signals including DNase and POLR2A (M), and cCRE activity quantified by MPRA experiments (N). *P*-values were calculated using Student’s *t*-test (* *P* < 0.05; ** *P* < 0.01; *** *P* < 0.001). (O) Enrichment of K562 regulatory variants in hub versus non-hub cCREs. *P*-values were calculated using binomial test (* *P* < 0.05; ** *P* < 0.01; *** *P* < 0.001).

Applying the transformation pipeline to the reconstructed cis-regulatory interaction matrices, we built 11,596 cis-regulatory networks for protein-coding genes in the autosomes of K562. We observed that the networks exhibit markedly different degrees of complexity, ranging from the simplest networks containing an isolated promoter to the most complex network comprising 93 tightly interconnected regulatory elements (*PFAS*, Fig. 3B). Therefore, we classified these cis-regulatory networks into simple, sparse, and complex modes based on network size and edges (Fig. 3B), and found that network complexity reflects distinct regulatory demands, even for members of the same protein family. For example, among STAT family members in K562, *STAT6* was governed by a simple network consisted of a promoter connected to less than 3 nearby CREs; *STAT2* and *STAT3* displayed sparse networks with more CREs but limited connectivity; whereas *STAT5A* and *STAT5B* were regulated by complex networks with dense interactions (Fig. 3C). Given the central role of *STAT5* in sustaining leukemic survival and proliferation^35^, the complex architectures of *STAT5A* and *STAT5B* suggest that disease-relevant STAT members require coordinated cis-regulatory control in K562. More broadly, these results indicate that functional divergence within a protein family^36^ may be accompanied by divergence in cis-regulatory architecture.

Extending the analysis to all K562 cis-regulatory networks further showed that the increased network complexity was accompanied by coordinated changes in topological properties, including enhanced transitivity and elevated average clustering coefficient (Fig. 3D). These features indicate dense communities or “small-world” properties emerging in the complex network, suggesting that complex networks are structured for hierarchical regulatory control. These unique properties of the network functionally manifest as genes regulated by complex cis-regulatory networks exhibiting significantly higher expression levels than those regulated by simple or sparse networks (Fig. 3E). Moreover, genes under the regulation of complex networks are preferentially enriched for cell identity markers, chronic myeloid leukemia-associated disease genes, while their constituent CREs showed higher enrichment in GWAS SNPs associated with blood traits (Fig. 3F).

### Modular organization supports transcriptional robustness in complex regulatory systems

Beyond global increases in network density, complex regulatory networks showed a substantial rise in indirect CRE-promoter interactions, in which regulatory influence is propagated through intermediate CREs rather than direct contact with promoter (Fig. 3G). This architecture implies a hierarchical regulatory logic, where information from distal elements is integrated through intermediate regulatory hubs, potentially buffering robust transcription against local perturbations.

This observation prompted further structural analysis, revealing that increased indirect CRE-promoter interactions result in modular organized structure of networks (Fig. 3H). Among all cis-regulatory networks of K562, 33.6% exhibit modular structures (Fig. 3I), and this proportion reaches as high as 96.1% in complex cis-regulatory networks (Fig. 3J, Supplementary Fig. S4A-C). In modular networks, CREs are partitioned into discrete modules characterized by dense intra-module interactions and sparser inter-module connections, often coordinated through a few functionally important hub CREs (Fig. 3H).

Notably, genes regulated by modular networks exhibited higher expression levels and lower Tau scores (namely lower tissue specificity) (Fig. 3K), reflecting an essential requirement for maintaining broad and high-level expression across tissues. Meanwhile, higher phastCons100way ^37^ scores of CREs that constitute modular network indicates stronger evolutionary conservation and functional constraint in these CREs (Fig. 3K). These features collectively demonstrate that hierarchical regulatory architectures are preferentially employed for genes requiring robust and stable transcription.

Within modular networks, hub CREs, defined by high within-module centrality and inter-module connectivity, emerged as key connectors between regulatory modules (Supplementary Fig. S5). Compared with non-hub CREs, higher degree centrality, betweenness centrality (Fig. 3L), and closeness centrality (Supplementary Fig. S6) of hub CREs confirm their role as integrators of regulatory information. Consistently, their elevated chromatin accessibility, RNA polymerase II occupancy (Fig. 3M), and other activation-associated epigenetic modification (Supplementary Fig. S7), as well as higher transcription activity quantified by MPRA experiment ^23^ (Fig. 3N) indicate their active involvement in transcriptional processes within K562. A recent study measured the regulatory activity of 221,412 trait-associated variants by quantifying their ability to alter transcriptional output in K562 ^38^. These experiment verified K562 regulatory variant are significantly enriched in hub CREs (Fig. 3O), suggesting that perturbation of these hubs may disproportionately disrupt inter-module coordination and transcriptional output. Collectively, these results indicate that regulatory network complexity is not a passive consequence of CRE abundance but reflects an hierarchical organization that enhances regulatory robustness and enables coordinated control through hub CREs ^39^.

### Modularity specificity acts as a topological signature of cell identity

Building on the observation that modular topology of regulatory network provides transcriptional robustness in K562, we next asked whether the deployment, and conversely, absence, of such architecture encodes specific functional requirements of a given cell type or disease state. To test this hypothesis, we reconstructed the cis-regulatory networks for all genes across nine cell lines. Through cross-cell line comparisons, we identified genes possessing modular cis-regulatory networks only in K562 (Specific modular genes), genes displaying non-modular networks only in K562 (Specific non-modular genes), genes possessing modular networks across all cell lines (Common modular gene), and genes possessing non-modular networks across all cell lines (Common non-modular gene). Utilizing these four special gene sets, we first investigated whether the specificity of cis-regulatory network modularity is associated with cell-type-specific gene regulation. In K562, specific modular genes were significantly enriched in K562-specific up-regulated genes, whereas specific non-modular genes preferentially overlapped with K562-specific down-regulated genes (Fig. 4A). This asymmetric association suggests that the emergence or disappearance of modular regulatory organization is not merely a structural property, but is functionally coupled to active or repressive transcriptional programs defining cell identity.

**Fig. 4.**
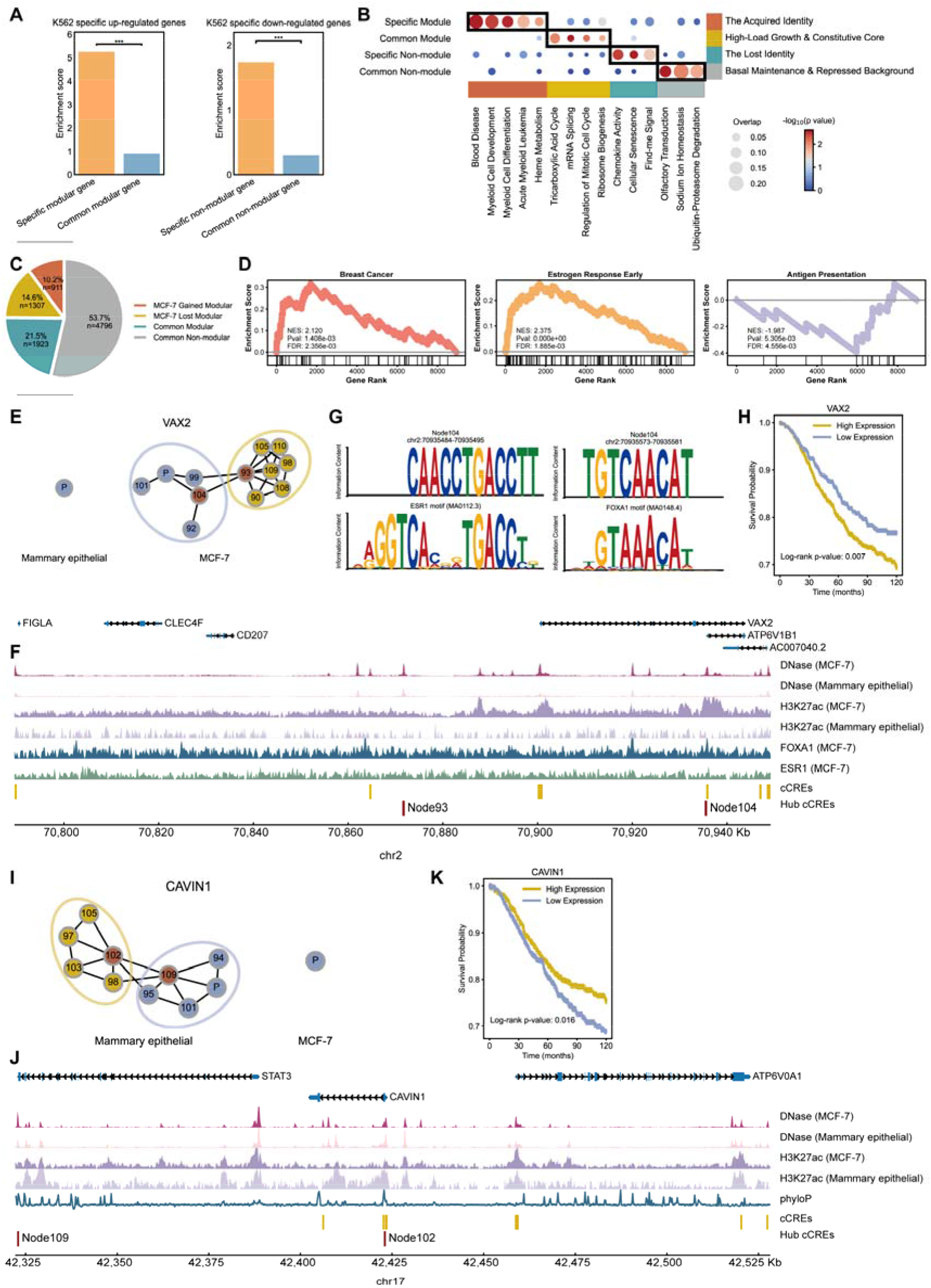
Cis-regulatory network modularity encodes cell identity and undergoes disease-specific rewiring. (A) Left: Enrichment of K562-specific up-regulated genes in specific versus common modular genes. Right: Enrichment of K562-specific down-regulated genes in specific versus common non-modular genes. *P*-values were calculated using binomial test (* *P* < 0.05; ** *P* < 0.01; *** *P* < 0.001). (B) Functional enrichment analysis of genes with specific modular structure, common modular structure, specific non-modular structure, and common non-modular structure in K562. The size of the bubbles represents the fraction of genes matched to the pathways, and the color indicates the statistical significance (-log10 *P*-value). Functional categories of genes with different types of modular structure are summarized on the right (e.g., “The Acquired Identity” for K562-specific modular gene). The color blocks sharing the same color on the right and bottom bridge gene functional categories to their corresponding pathways. (C) Pie plot showing the percentage of cis-regulatory networks that gained modularity, lost modularity, or remained as common modular/non-modular networks in MCF-7 compared to mammary epithelial cell. (D) Gene Set Enrichment Analysis (GSEA) reveals enriched pathways identified using network change scores between MCF-7 and mammary epithelial cell networks. (E-H) Analysis of *VAX2* gene, a representative of “MCF-7 gained modular” genes. (E) *VAX2* shows a complex, modular network in MCF-7 (right) compared to a simple, non-modular structure in mammary epithelial cell (left). (F) Genome browser view of the *VAX2* locus displaying tracks for DNase-seq, H3K27ac ChIP-seq, and transcription factor (FOXA1, ESR1) binding in MCF-7 and mammary epithelial cell. The gained modular network in MCF-7 corresponds to increased active epigenetic signals and TF binding at network CREs, especially hub CREs. (G) Motif analysis confirms the presence of ESR1 and FOXA1 binding motifs within the sequence of the identified hub cCRE, Node104. (H) Kaplan-Meier survival curve shows that high expression of *VAX2* is significantly associated with shorter survival in breast cancer patients. (I-K) Analysis of *CAVIN1* gene, a representative of “MCF-7 lost modular” genes. (I) *CAVIN1* shows a simple, non-modular network in MCF-7 (right) compared to a complex, modular structure in mammary epithelial cell (left). (J) Genome browser view of the *CAVIN1* locus displaying DNase-seq, H3K27ac ChIP-seq, and phyloP tracks. The loss of the modularity in MCF-7 corresponds to a decrease in epigenetic signals at CREs. (H) Kaplan-Meier survival curve shows that high expression of *CAVIN1* is significantly associated with better overall survival in breast cancer patients.

Therefore, we hypothesized that cell line-specific gene programs can be reflected through modular regulatory architectures, while non-modular networks may support transcriptional repression or basal regulation, and that transitions between these architectures reflect regulatory rewiring during different states. Consistent with this notion, functional enrichment analysis ^40^ revealed a clear partition of biological roles across modularity categories (Fig. 4B). K562-specific modular genes were enriched for pathways underlying hematopoietic lineage and oncogenic drivers associated with leukemia, representing an “acquired identity” unique to the lineage-specific and malignant state of K562. Conversely, K562-specific non-modular genes were enriched for pathways underlying immune response and cellular senescence, reflecting the contextual shutdown of relevant transcriptional programs to escape survival pressures, representing a “lost identity”. We further observed that conserved modularity patterns were associated with the consistent survival requirements across multiple cell types. For example, genes with common modular structures across diverse cell types (common module) were those ubiquitously high-expressed genes. The preservation of modular structure for these genes implies the need for complex regulation to support rapid cellular proliferation and high metabolic burden. As a result, these genes are enriched for high-demand basic metabolic and biosynthetic processes. Meanwhile, genes with common non-modular structures across diverse cell types (common non-module) were those ubiquitously low-expressed genes. This group consists of housekeeping genes responsible for fundamental homeostasis and remaining silent genes, indicating that simple regulatory architectures may suffice for genes with limited regulatory demands (Fig. 4B).

### State-dependent rewiring of regulatory topology captures disease-associated reprogramming

After confirming the association between modularity specificity and cell identity, we further exploited this characteristic to examine how network modularity is rewired during disease progression. Specifically, we systematically analyzed the gain and loss of network modularity during oncogenic transformation from mammary epithelial to MCF-7. A substantial fraction of cis-regulatory network gained (10.2%) or lost (14.6%) modular organization during malignant transformation (Fig. 4C).

To quantify the magnitude of network dynamics between disease and healthy states, we defined a network dynamic score Scor*e*_dynamic_ which measures global changes in interaction strength. Genes undergo regulatory complexity expansion have Scor*e*_dynamic_ larger than zero, and vice versa. Gene set enrichment analysis (GSEA) ^41^ based on network dynamic scores revealed that genes gaining complexity in MCF-7 were strongly associated with breast cancer-related pathways, including estrogen response and oncogenic signaling, whereas genes losing complexity were associated with antigen presentation pathway, suggesting a collapse of CREs coordination that leads to transcriptional silencing of immune-related programs during tumor progression (Fig. 4D). These results suggest that disease-associated regulatory reprogramming is accompanied by systematic remodeling of network hierarchy.

*VAX2* gene exemplifies modularity gain in MCF-7. While *VAX2* is regulated by a simple, non-modular network in mammary epithelial cell, it acquires a complex, modular regulatory architecture in MCF-7 (Fig. 4E). This structural transition coincides with increased chromatin accessibility and H3K27ac signals, as well as enhanced binding of breast cancer-specific pioneer transcription factors, ESR1 (ERα) and FOXA1, at network CREs, particularly at hub CREs (node 93, EH38E3351126 & node 104, EH38E3351172) (Fig. 4F). Motif analysis further confirmed the presence of ESR1 and FOXA1 motifs within hub CRE 104 (Fig. 4G). ESR1 and FOXA1 cooperatively open chromatin and establish estrogen-responsive cis-regulatory programs in breast cancer ^42,43^. In this context, hub CRE that bound by ESR1/FOXA1 may act as an integrator to amplify and stabilize estrogen-driven regulatory information, thereby promoting aberrant activation of *VAX2*. Recently, a study identified a high-risk mesenchymal development subtype of breast cancer, where the *VAX2* is the master regulator of this subtype ^44^. The following experiment confirms that the elevated *VAX2* expression is driven by a breast cancer-specific super enhancer, which coincides with the MCF-7 gained modularity of cis-regulatory network of *VAX2* identified by ORIGAMI. Clinically, elevated *VAX2* expression is associated with poorer survival in breast cancer patients (Fig. 4H), attributing oncogenic activation and adverse outcomes to the acquired cis-regulatory modularity of *VAX2*.

Conversely, *CAVIN1* (*PTRF*) experienced loss of modularity during tumorigenesis. CAVIN1 is a core structural component of caveolae that regulates membrane signaling and lipid homeostasis, and its loss is frequently associated with enhanced tumor progression due to disrupted signal compartmentalization ^45,46^. In mammary epithelial cell, *CAVIN1* is regulated by a complex modular network, which collapses into a simpler, non-modular structure in MCF-7 (Fig. 4I). This loss of hierarchy may induced by reduced chromatin accessibility and activity across network CREs, particularly at evolutionarily conserved hub CREs (node 102, EH38E3224086 & node 109, EH38E3223986) marked by high phyloP scores ^47^ (Fig. 4J). Consistent with the collapse of network, the expression of *CAVIN1* is down-regulated in MCF-7, and lower expression is associated with shorter patient survival (Fig. 4K), suggesting that the disruption of well structured cis-regulation in healthy state may contribute to silencing of tumor-suppressive programs. Together, these analyses demonstrate that rather than being a static property, the dynamic of network modularity reflects disease-specific reconfiguration of transcriptional control ^48–50^.

### Inter-module interactions generate synergistic control in hierarchical regulatory networks

We next examined how multiple modules collaboratively contribute to gene regulation, beginning with the difference in the regulatory properties of intra- and inter-module interactions. CRE pairs within the same module are generally closer in linear genome distance (Fig. 5A) and exhibited higher chromatin co-accessibility (Fig. 5B) than those across modules. Additionally, Pol II-mediated ChIA-PET ^3^ verified that intra-module CRE pairs are physically more proximate, forming compact within-module interactions (Fig. 5C). To investigate how TFs involve in gene regulation, we quantified the TF motif similarity between CRE pairs across three groups using cosine similarity. The result shown that CRE pairs within the same module display generally higher similarity in TF motifs (Fig. 5D), suggest that CREs within same module may share similar TF binding profiles.

**Fig. 5.**
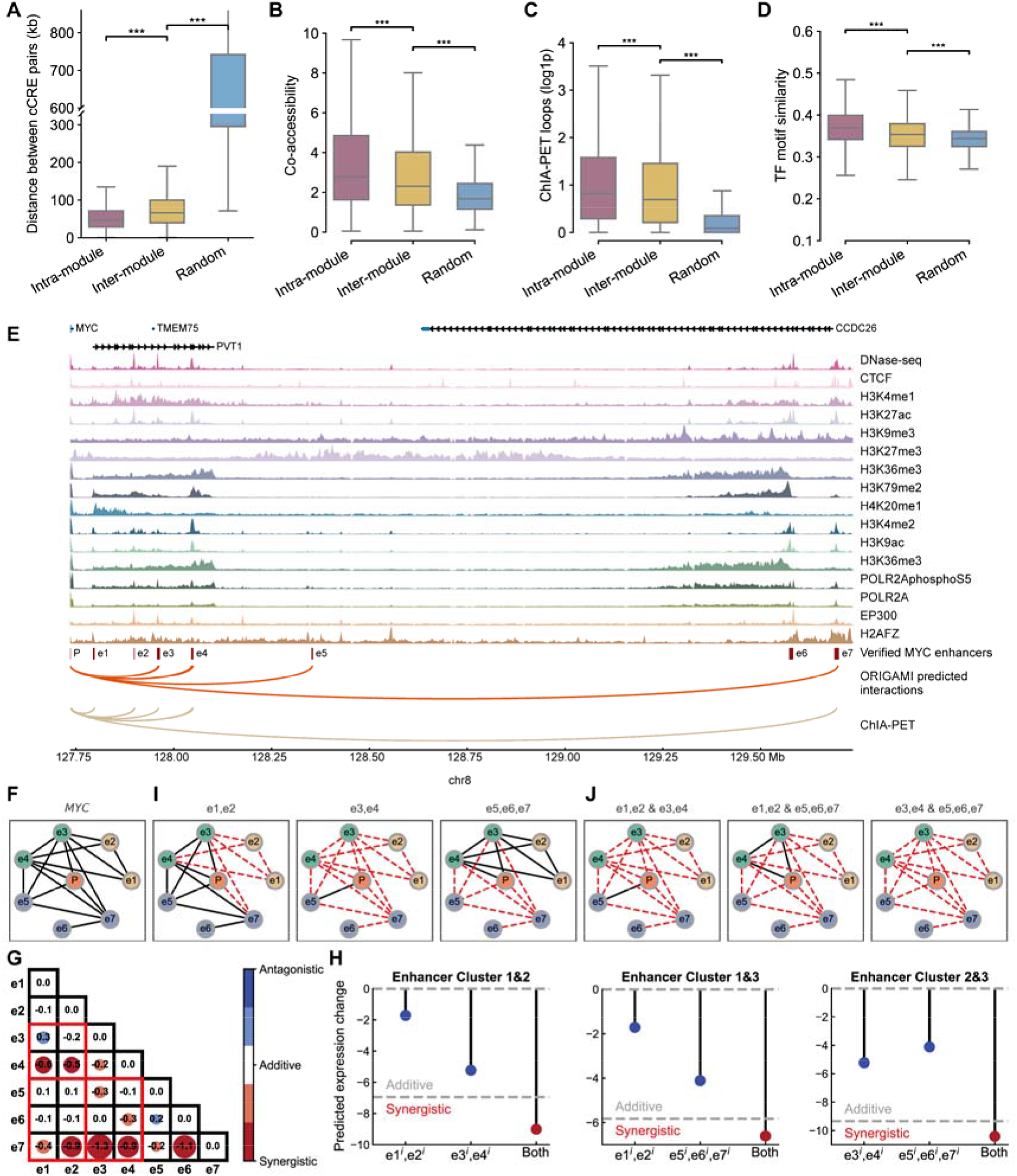
Synergistic transcriptional control emerges from inter-module cis-regulatory interactions at the *MYC* locus. (A-D) Comparison between intra-module, inter-module and random CRE pairs in terms of genomic distance (A), co-accessibility (B), ChIA-PET interaction frequency (C), and transcription factor (TF) motif similarity (D). *P*-values were calculated using Student’s *t*-test (*** *P* < 0.001). (E) Genome browser view of the *MYC* locus showing epigenomic tracks, experimentally verified *MYC* enhancers (e1-e7), and CRE-promoter interactions predicted by ORIGAMI compared to ChIA-PET interactions in K562. (F) The reconstructed *MYC* cis-regulatory network. (G-H) Analysis of combinatorial effects of enhancer knockouts. At the enhancer level (G), the values in the heatmap indicate the predicted expression change from combined enhancer knockouts compared to the additive of individual knockouts of two enhancer. And the regulatory effects between enhancer pairs are classified as synergistic (red), additive (white), or antagonistic (blue) according to the difference of predicted expression change. At the enhancer cluster level (H), lollipop plots show the predicted expression change from combined enhancer cluster knockouts (red dots) compared to the expected additive effect (grey dashed line), confirming synergistic interactions between inter-module enhancer clusters. (I-J) The perturbed networks after in silico knockouts of individual (I) and combinations (J) of enhancer clusters. Red dashed lines indicating lost interactions.

A recent study has experimentally confirmed seven active enhancers in K562 ^18^, spanning the genome sequence to as far as 2 Mb downstream of *MYC* TSS, and a following study revealed a nested epistasis cis-regulatory network formed by these enhancers ^51^. Inspired by these findings, we first examined if ORIGAMI could restore the nested cis-regulatory network of *MYC* accurately. Then, based on the reconstructed network, we evaluated whether perturbations on inter-module enhancer pairs could alter gene regulation in a synergistic manner. As shown in Fig. 5E, ORIGAMI accurately recovered CRE-promoter interactions supported by Pol II-mediated ChIA-PET ^3^, especially ultra long-range interactions between *MYC* promoter (P) and enhancer 7 (e7). The reconstructed *MYC* cis-regulatory network, shown in Fig. 5F, further demonstrates that the long-range enhancer 6 (e6) participates in gene regulation by indirectly interacting with the promoter through e7.

Then, to quantitatively assess whether enhancer interactions act additively or synergistically, we compared the predicted gene expression change resulting from combined enhancer knockouts with the expected additive effect derived from individual knockouts. For each enhancer pair (*e*_*i*_, *e*_*j*_), we computed the difference between the expression change predicted for the joint knockout and the sum of expression changes predicted for the corresponding single knockouts. Negative deviations indicate synergistic interactions, where joint perturbations produce a stronger-than-expected repression of expression, whereas positive deviations indicate antagonistic effects. Three modules were detected in the *MYC* network, namely e1&e2, e3&e4, and e5-e7. Fig. 5G summarizes this analysis across enhancer pairs, in which inter-module enhancer pairs (in the red frame) revealed widespread synergistic regulatory effects with negative deviations (red), while intra-module pairs close to zero, consistent with additive or redundant regulation within modules. Further analysis at the enhancer cluster level also confirmed the same effect, where the predicted expression change from combined knockouts of enhancer clusters (red dots) consistently exceeds the additive expectation (grey dashed line) (Fig. 5H). The changes in network structure also shown that while knockout of single enhancer clusters led to partial attenuation of network (Fig. 5I), simultaneous disruption from different modules caused disproportionately collapse of whole network (Fig. 5J). Together, these analyses demonstrate that inter-module interactions encode synergistic regulatory logic, providing functional evidence that modular cis-regulatory architecture enables non-linear integration of distal regulatory signals.

### In silico perturbations link local regulatory disruption to network-scale response

To interrogate whether ORIGAMI could accurately predict perturbation effects and how variants perturb cis-regulatory networks, we performed systematic in silico perturbations targeting both cis-acting sequence variants and trans-acting TF motifs using ORIGAMI.

We first assessed cis-regulatory effects by introducing fine-mapped eQTL variants of breast mammary ^52^ into the input genome sequences and predicting the resulting expression changes. ORIGAMI correctly recapitulated the directionality of eQTL effects, with predicted expression changes significantly shifted above or below zero for up- and down-regulated eQTLs, respectively (Fig. 6A). Importantly, the magnitude of predicted expression changes positively correlated with GTEx-derived eQTL effect sizes (Supplementary Fig. S9). Especially for the eQTLs near TSS, we observed a positive correlation with a Spearman ρ of□□0.56 (Fig. 6B), indicating that ORIGAMI captures quantitative cis-regulatory impacts. Attribution analysis further revealed the mechanistic basis of these effects: for a representative variant rs760077, which caused significant expression down-regulation of *THBS3*, the mutation disrupted BATF3 binding motif in a regulatory element of *THBS3*, producing a localized loss of attribution signal and reduced gene expression (Fig. 6C). These results demonstrate that ORIGAMI links nucleotide-level sequence perturbations to functional transcriptional consequences by modeling motif-level regulatory disruption.

**Fig. 6.**
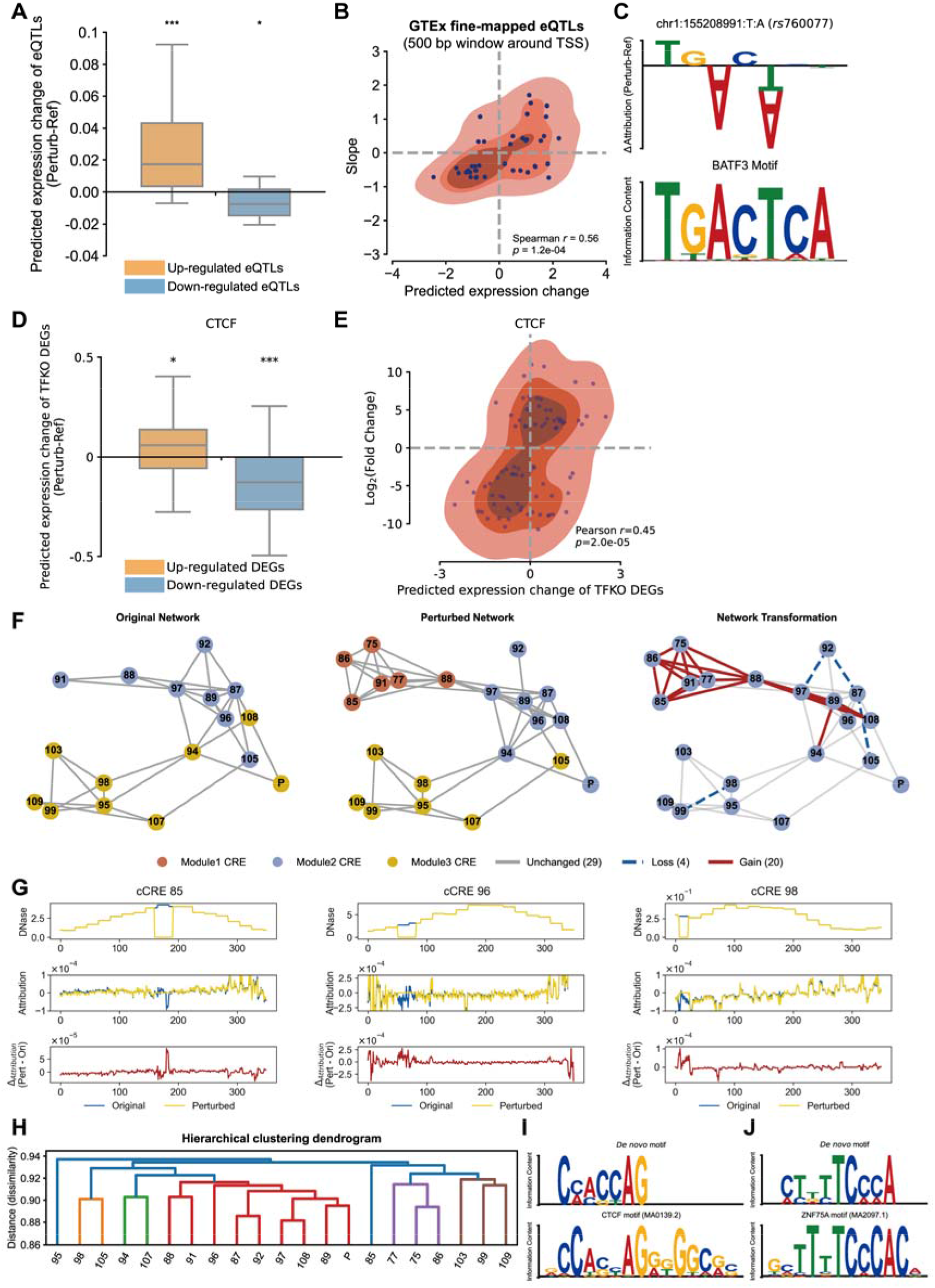
in silico perturbation analysis of cis- and trans-regulatory variants. (A) Boxplot showing the distribution of ORIGAMI-predicted expression changes after in silico eQTL mutation within a 500 bp window around the TSS in mammary epithelial cell. eQTLs are categorized as up-regulated or down-regulated based on their effect size. *P*-values were computed through one-sided Mann-Whitney U test to evaluate if the predicted expression change were significantly larger or smaller than 0, respectively for the up- or down-regulated eQTLs (* *P* < 0.05; *** *P* < 0.001). (B) The correlation between the predicted expression changes and the GTEx derived slopes for fine-mapped eQTLs within a 500 bp window around the TSS. (C) Attribution analysis of a representative eQTL (rs760077). The top motif logo illustrates the attribution change (ΔAttribution) caused by the T to A variant, which disrupts the BATF3 binding motif shown below and results in decreased gene expression. (D) Boxplot showing the distribution of ORIGAMI-predicted expression changes for up- and down-regulated DEGs after in silico CTCF knockout in K562. *P*-values were computed through one-sided Mann-Whitney U test to evaluate if the predicted expression change were significantly larger or smaller than 0, respectively for the up- or down-regulated DEGs (* *P* < 0.05; *** *P* < 0.001). (E) The correlation between the predicted expression changes and the log2(fold change) derived from CTCF knockout experiment. (F) Visualization of the cis-regulatory network for a representative gene *PROCR* before (Original) and after (Perturbed) in silico CTCF knockout in K562. The network transformation highlights interactions that are unchanged (grey), lost (blue dashed), or gained (red solid) after to the perturbation. (G) Epigenetic-based attribution analysis of three perturbed CREs, which contain CTCF motif, from the network in (F). Tracks show the DNase signal, attribution of DNase signal, and attribution change (ΔAttribution) for the original and perturbed states, revealing nucleotide-resolution changes in regulatory potential. (H) Hierarchical clustering dendrogram of the CREs in the *PROCR* cis-regulatory network. Cluster labeled in same color represent a group of CREs that share similar interaction changes after CTCF knockout. (I-J) DNA attribution analysis of CREs in the network (F). (I) The *de novo* motif identified from genomic segments with the largest attribution change in each CRE (top) matches known CTCF binding sites (bottom). (J) The *de novo* motif identified from genomic segments with the largest attribution after CTCF knockout in each CRE (top) matches known ZNF75A binding sites (bottom).

We next examined trans-regulatory perturbations by simulating TF knockouts in K562 through down-regulating activation-associated epigenetic signals and up-regulating repression-associated epigenetic signals on detected TF motifs in the CREs. Across 34 TF knockout experiments ^3,53^, ORIGAMI’s predicted expression changes showed consistency with experimental perturbation outcomes of 22 TF knockouts, with the predicted direction and magnitude of expression changes for both up- and down-regulated genes correctly matched the experimental results (Supplementary Fig. S10). Taken CTCF knockout as an example, ORIGAMI predicted significant expression decreases for down-regulated genes and increases for up-regulated genes following CTCF loss (Fig. 6D), where 74% down-regulated genes with a predicted expression change less than 0, and 64% up-regulated genes with a predicted expression change larger than 0. Moreover, the Pearson correlation coefficient between the predicted and observed log2 fold changes of DEGs is 0.45 (Fig. 6E), indicating that the model reliably captures magnitudes of TF regulatory effects.

Then, we utilized the cis-regulatory network rewiring and expression change of the gene *PROCR* following CTCF knockout to jointly elucidate how ORIGAMI bridges trans-regulatory disruption with downstream effects. After CTCF knockout, the expression of *PROCR* significantly increased, accompanied by markedly enhanced complexity and modularity within its cis-regulatory network. Specifically, the perturbed network gained four new CREs, forming a novel regulatory module tightly connected by twenty edges (red edges) (Fig. 6F). The dynamic cluster detection (the adjacency difference matrix **D** and dissimilarity matrix **T** of *PROCR* are shown in Supplementary Fig. S11 and Supplementary Fig. S12, respectively) also revealed a covariant clusters, composed of nine nodes within Module 1 (cluster marked in red) (Fig. 6H), which means that these nodes exhibited similar connectivity alterations after perturbation.

Next, we performed gradient × input analysis of DNase signals. Figure 6G showed the results on three CREs which contain CTCF motif. The results indicate that the original open state of the genomic regions containing CTCF motif contributed negatively to gene expression. Therefore, after simulating the closure of these regions, the model predicted an increase in gene expression level. Analysis of attributions on genome sequence also discovered a *de novo* motif enriched in genomic segments with the largest differential attribution (Fig. 6I, upper panel). Using TomTom ^54^ to compare this *de novo* motif against known motifs in JASPAR database ^55^, we found it matches the CTCF motif (Fig. 6I, lower panel), indicating the model captures the significant regulatory impact of CTCF binding sites following knockouts. To further investigate which TF drives the upregulation of PROCR expression following CTCF binding disruption, we analyzed attribution profiles of genome sequence after perturbation. Among the most contributory genomic segments, we detected a *de novo* motif matching the motif of transcription factor ZNF75A (Fig. 6J), suggesting that ZNF75A may dominate the expression regulation of *PROCR* after CTCF knockout as a compensatory or secondary regulatory mechanism. Together, these analyses demonstrate that ORIGAMI not only predicts the transcriptional consequences of cis- and trans-regulatory perturbations, but also resolves the underlying network-level rewiring and motif-specific mechanisms through which regulatory variants reshape gene expression programs.

## Discussion

A central challenge in regulatory genomics is to bridge static, noisy measurements of chromatin state with the dynamic, causal control of gene expression ^25,39,49,50,53,56^. Here, we formalize this problem as a latent graph inference task. By constraining network reconstruction with gene expression prediction, ORIGAMI learns regulatory architectures that capture both high-order genome organization and functional causality, rather than structural proximity alone.

One key insight emerging from this work is that regulatory topological regimes is not uniformly distributed across the genes, but instead reflects tissue-specific functional demands. The hierarchically organized, modular regulatory architectures govern robust and highly expressed genes, whereas simpler, non-modular networks are associated with fundamental and low-cost regulation. Within these regimes, regulatory modules do not act independently; instead, hub CREs mediate inter-module coupling, enabling non-linear integration of regulatory signals ^12,29,39,48,50^. Therefore, such hierarchical topology confers both robustness against local perturbations and flexibility for coordinated regulatory responses.

The dynamic rewiring of regulatory modularity across cell states further underscores modular topology as a regulatory strategy rather than a static feature ^15,57^. In this context, tumorigenesis can be viewed as a systematic topological transition of cis-regulatory networks, which is characterized by network-scale rewiring dynamics, rather than isolated molecular alterations. Such transitions reshape the flow of regulatory information, amplifying oncogenic programs while attenuating tumor-suppressive pathways. These observations imply that pathological states may exploit regulatory rewiring to achieve stable yet aberrant expression programs, offering a new perspective on how regulatory networks are repurposed during tumorigenesis.

A particularly notable aspect of ORIGAMI is its ability to accurately predict the transcriptional effects of both cis- and trans-regulatory perturbations, positioning the model as a simulation-ready regulatory system. This capability aligns closely with the emerging paradigm of virtual cell modeling ^58–60^, which enable systematic, in silico interrogation of genetic perturbations without experimental manipulation. By linking genetic and epigenetic variations to network topology and transcriptional outcomes, ORIGAMI functions as a virtual cell engine, demonstrating how virtual perturbations can be used to prioritize functional variants, explore regulatory causality, and discover disease mechanism arising from noncoding variations.

Despite these advances, several limitations should be acknowledged. First, ORIGAMI infers regulatory interactions from population-averaged bulk data, and therefore does not explicitly capture continuous cell state variation. Extending this framework to single-cell multi-omics data may enable mapping of continuous regulatory state-space trajectories and improve prediction of heterogeneous perturbation responses within microenvironments. Second, while ORIGAMI reconstructs gene-centric cis-regulatory interactions, it does not explicitly model feedback regulation or trans-regulatory coupling. Integrating gene-centric networks into gene regulatory networks that incorporate trans-regulatory information may enable simulation of full-scale cellular dynamics, advancing toward more comprehensive and authentic regulatory circuits.

Looking forward, ORIGAMI establishes a generalizable computational framework for modeling gene regulation as a structured, multi-scale, and perturbable system, which provides a foundation for predictive modeling of regulatory variation and disease-associated network reprogramming. Future work may extend this framework for variant interpretation, disease modeling, and the rational design of interested perturbation experiments. More broadly, it advances the development of virtual cell models that enable mechanistic and quantitative exploration of cellular regulation in silico.

## Methods

### Model architecture

We formulate cis-regulation as a latent graph inference problem under multi-omics and noisy observations, where transcriptional output provides a functional constraint on graph reconstruction. For each target gene **g**, ORIGAMI considered a set of (*N =* 200) cCREs located up- and down-stream of the transcription start site of **g**, together with a promoter of **g** itself. Each cCRE and the promoter are represented at base-level using multi-omics data, including nucleotide sequence in the form of one-hot encoding, DNase-seq signals in the form of read-depth normalized signal and Histone/TF ChIP-seq signals in the form of fold change over control (Supplementary Table S1). Chromatin conformation data derived from Hi-C provides a sparse and noisy prior over potential regulatory interactions between cCREs.

The objective of model is to (i) infer a reconstructed cis-regulatory interaction matrix 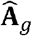 among cCREs and the promoter, and (ii) predict the gene expression value *ŷ*_*g*_.

The backbone of model consists of three modules: (i) a hierarchical transformer module to extract regulatory features of regulatory elements; (ii) a graph autoencoder module to reconstruct regulatory network; and (iii) a graph regression module to predict gene expression.

### Hierarchical Transformer for cCRE Regulatory Representation

1. Base-Level Embedding The nucleotide sequence (S_*i*_) and epigenomic tracks (E_*i*_) of the *i*-th cCRE are mapped to learnable base-level embeddings through a linear projection, respectively. The model then sums the outputs from the two embedding layers, as:

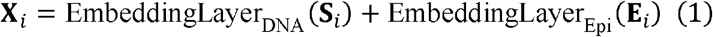

yielding a base-level representation **X**_*i*_ ∈ ℝ^*L*× *d*^ of cCR*E*_*i*_.
2. CRE-centric Transformer Encoder To aggregate the base-level representation into a single cCRE representation, we concatenate a learnable [CLS] token to **X**_i_ and apply a CRE-centric Transformer Encoder:

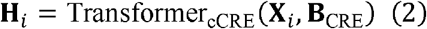

where **B**_CRE_ encodes bucketed relative position bias ^61^. This bias encodes pairwise relative distances between base positions, discretized into logarithmically scaled buckets. This design introduces the positional information and allows the model to retain sensitivity to local motif spacing and short-range regulatory grammar. Formally, for positions *i* and *j* within a cCRE, the position bias is defined as:

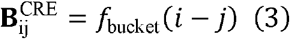

where *f*_bucket_ (·) maps relative distances to a fixed number of learnable bias buckets, shared across attention heads. During training, the [CLS] token can adaptively attend across bases and learn a holistic cCRE representation **h**_*i*_.

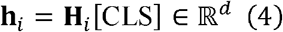

Specifically, the CRE-centric Transformer Encoder consists of 2 stacked position-aware transformer blocks with 4 heads. For attention head *h*, queries **Q**_*h*_, keys **K**_*h*_, and values **V**_*h*_ are computed as:

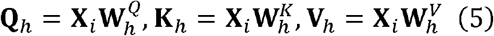

where 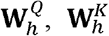 and 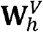 are trainable weight matrices ^62^. The position-aware multi-head attention output is:

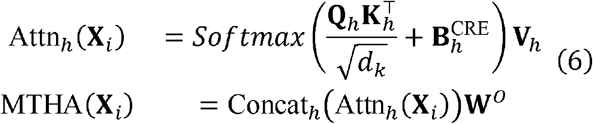

where *d*_*k*_ denotes the dimension of the Key vector and **W**^*o*^ denotes a learnable matrix used for linearly transform the concatenated outputs from all attention heads.
3. Regulatory-centric Transformer Encoder To capture higher-order interactions among regulatory elements, all cCRE embeddings and the promoter embedding are concatenated:

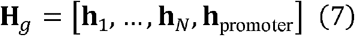

**H**_*g*_ is subsequently fed into a Regulatory-centric Transformer Encoder, which shares similar architecture with CRE-centric Transformer Encoder.

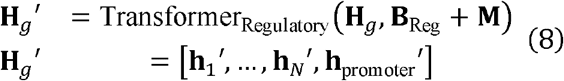

Here, **M** is an attention mask enforcing unidirectional information flow from cCRE to promoter. **B**_Reg_ is a long-range relative position bias designed to capture long-range and heterogeneous genomic distances, spanning several orders of magnitude. Given genomic coordinates *p*_*i*_ and *p*_*j*_ of regulatory elements *i* and *j*, the bias is computed as a sum of relative position embeddings evaluated at different distance scales:

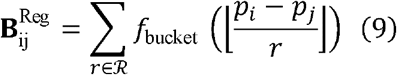

where ℛ = {150,600,2400,10,000,40,000} represents a set of predefined genomic resolutions (in base pairs). Each scale contributes an independent learnable bias term, and the resulting biases are summed across scales and attention heads. Finally, the output **H**_*g*′_ ∈ ℝ ^(*N*+1)×d^ provides regulatory-aware embeddings for **h**′ all regulatory elements of target gene **g**.

### Graph Autoencoder for Regulatory Network Reconstruction

To explicitly model hierarchical interactions among regulatory elements, we formulate the regulatory landscape of each target gene as a graph and reconstruct its latent regulatory network using a Graph Autoencoder.

1. Graph Construction For each gene **g**, we define a graph *G*_*g*_ = (*V*_*g*_, ℰ_*g*_). Nodes *V*_*g*_ = {*v*_1_, … *v*_*N*_, *v*_promoter_} correspond to the *N* cCREs and the promoter. The regulatory-aware embeddings **h**′ are used as initial node features. Edges ℰ_*g*_ are initialized using Hi-C-derived chromatin contacts, providing a sparse and noisy structural prior over potential regulatory interactions.
2. Graph Attention Network (GAT) Encoder To encode contextual dependencies between regulatory elements, we apply a GAT ^63^ encoder to learn latent embedding **z**_*i*_ for node *v*_*i*_ :

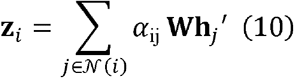

where *N*(*i*) denotes the neighbors of node *v*_*i*_, and the attention coefficient α_ij_ is defined as:

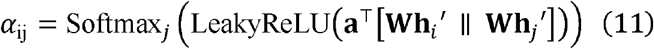

where **a**,**W** are learned, ‖ denotes vector concatenation, and Softmax_*j*_(·) normalizes the attention scores across all neighbors *j* ∈ *N* (*i*) This formulation enables the model to adaptively weight regulatory interactions, allowing influential cCREs to exert stronger effects on network topology.
3. Euclidean Distance-Based Decoder To reconstruct a denoised cis-regulatory interaction matrix 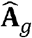, we decode edge probabilities using an euclidean distance-based decoder:

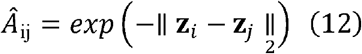

where *Â*_ij_ ∈ (0,1] represents the predicted interaction probability between regulatory elements *i* and *j*.

### Graph-Level Regression for Gene Expression Prediction

Not all reconstructed interactions in 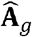 are functionally meaningful. Therefore, to constrain the model toward biologically meaningful interactions, the reconstructed graph is constrained by a downstream gene expression prediction task.

1. Network Pruning and Binarization The reconstructed interaction matrix is 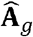 thresholded to retain high-confidence regulatory interactions:

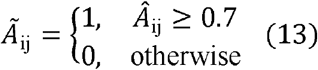

This step yields a sparse regulatory graph reflecting putative functional interactions.
2. Differentiable Graph Pooling (DiffPool) To derive a graph-level representation, we apply DiffPool ^64^ to hierarchically aggregate node embeddings:

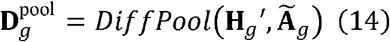

DiffPool is a deep, multi-layer GNN models, where GraphSAGE ^65^ is employed as the base GNN at every layer. At each layer, DiffPool learns differentiable soft assignment of nodes, mapping nodes to a set of clusters based on their learned embeddings and derived a coarsened graph. All nodes at the final layer are assigned to a single cluster, generating a final embedding vector 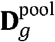 corresponding to the entire graph.
3. Gene Expression Regression The pooled graph representation is mapped to gene expression using a prediction head comprised of a linear projection and a soft plus activation:

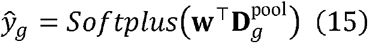

where *ŷ* denotes the predicted expression value of gene **g**.

### Learning Objective

The model is trained end-to-end by jointly optimizing a network reconstruction loss and a gene expression prediction loss. The graph autoencoder is supervised using a binary cross-entropy loss ℒ_recon_ on predicted and observed edges.

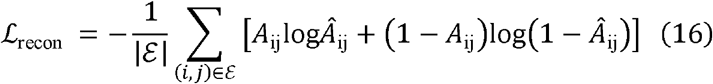

In parallel, gene expression is supervised using a Poisson negative log-likelihood loss ℒ_exp_ applied to log-transformed ground truth and predicted expression values.

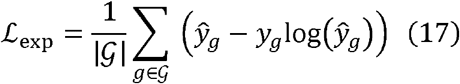

The overall objective is a weighted sum of the two terms,

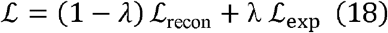

where *λ* = 0.3 balances network reconstruction and expression prediction tasks.

### Training data collection and preprocessing

Genome sequence and gene annotation were downloaded from GENCODE ^66,67^ (https://www.gencodegenes.org/human/release_30.html). RNA expression (in transcript quantifications format), DNase-seq (in read-depth normalized signal format), 15 types of Histone/TF ChIP-seq (in fold change over control format) and intact Hi-C (in pairs format) files across nine cell lines were downloaded from ENCODE ^2^ (https://www.encodeproject.org/). The ENCODE accessions for the downloaded files can be found in Supplementary Table S1. The cell type-specific cCREs were downloaded from SCREEN Registry V3^3^ (https://screen.encodeproject.org/#). DNase-only and low-DNase cCREs were removed. All cCREs were padded to 350 bp for the convenience of modeling.

When processing input data, we only considered protein-coding genes with annotations in GENCODE and RNA expression values. For each retained gene, we extracted 100 cCREs upstream (labeled as 0 to 99 from farthest to nearest distance from the TSS) and 100 cCREs downstream (labeled as 101 to 200 from nearest to farthest distance from the TSS) of its TSS, and padding the TSS to 350 bp as promoter, thus generated 201 regulatory elements for each gene. Based on the genomic coordinates of regulatory elements, we extracted the genome sequences, DNase-seq and Histone/TF ChIP-seq profiles of the corresponding regions. The genome sequences were one-hot encoded and the epigenome values were transformed using inverse hyperbolic sine (arcsinh) ^68,69^. The gene expression level was calculated as the sum of its transcripts TPM and Log1p transformed. The downloaded Hi-C data in pairs format were binned into 350bp-resolution Hi-C contact matrix in cooler format ^70^ (details in Supplementary Methods).

### Method evaluation

We benchmarked ORIGAMI’s expression prediction performance against four baselines that specialize in both gene regulation and expression modeling. These methods represent three distinct categories of modeling approaches: (1) pre-training and fine-tuning framework-based methods like EPCOT-CNN and EPCOT-LSTM^71^; (2) attention mechanism-based method like CREaTor^32^; and (3) graph neural network-based method like GraphReg^27^. Following instructions in their published literature and GitHub documentation, we retrained each method on training data that was as consistent as possible with ORIGAMI for a fair comparison (details in Supplementary Methods), especially preserving the omics information from chromosomes 8 and 9 of K562 as the zero-shot test set.

To evaluate the performance of cis-regulatory network reconstruction on K562, we compared ORIGAMI against six baseline methods: ABC contact^19^, Enformer^25^, GraphReg^27^, CREaTor^28^, cCRE contact matrix and 1/dist, on their ability to prioritize CRISPR-validated K562 cCRE-gene pairs (details in Supplementary Methods). All the methods were evaluated by the experimentally validated K562 enhancer-target gene pairs that exhibited a significant expression change after perturbation ^19,72,73^. The genomic coordinates of the enhancers were converted from hg19 to hg38 using liftover through UCSC Genome Browser ^74^ (http://genome.ucsc.edu/).

To evaluate whether the cis-regulatory matrices align with the 3D genome organization, we obtained the K562 TAD boundaries at base-pair resolution identified by preciseTAD ^34^ and chromatin loops from the ENCODE portal (ENCFF256ZMD).

### Network structure analysis

#### Transforming cis-regulatory interaction matrix into cis-regulatory network

First, the reconstructed cis-regulatory interaction matrix 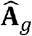 is thresholded to retain high-confidence regulatory interactions:

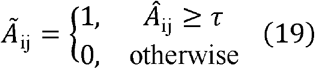

where *τ* is set to 0.9 by default unless otherwise specified below.

Next, we transformed the filtered cis-regulatory interaction matrix into a network structure using networkx ^75^. Finally, we retained 1-, 2- and 3-hop neighbors of promoter and all the edges between these nodes, thus derived a cis-regulatory network of target gene.

#### Network complexity classification and associated gene characteristic analysis

The cis-regulatory networks are classified into simple, sparse, and complex modes based on the number of network nodes and edges. Simple networks contain fewer than three nodes; sparse networks have more than three nodes but fewer than 100 edges; and all remaining networks are categorized as complex networks.

The cell type-specific identity marker were retrieved from CellMarker 2.0 ^76^ and PanglaoDB database ^77^. We use the cell types listed in Supplementary Table S3 and Supplementary Table S4 to filter marker in CellMarker 2.0 and PanglaoDB, respectively. The final filtered identity markers of K562 are listed in Supplementary Table S5. The chronic myeloid leukemia-associated disease genes were retrieved from four chronic myeloid leukemia pathways from KEGG ^78^, BioPlanet^79^, Elsevier pathway collection^80^, and DisGeNET^81^, respectively, which are listed in Supplementary Table S6. The GWAS Catalog SNPs were downloaded through the UCSC Genome Browser ^74^ (http://genome.ucsc.edu/), and we use the keys listed in Supplementary Table S7 to filter GWAS SNPs associated with blood traits.

#### Network module detection and associated gene characteristic analysis

We applied the Louvain Community Detection Algorithm for module detection using NetworkX ^75^. We used RNA expression for nine cell lines (listed in Supplementary Table S1) to calculate the tau scores for genes govern by modular and non-modular networks. RNA expression across multiple technical or biological replicates within the same cell line was averaged. The Tau score was calculated as

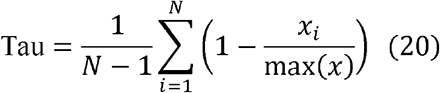

where *x*_*i*_ is the expression level of a gene in cell line *i* and *N* is the number of cell lines. Genes with a tau score close to 1 are more specifically expressed in one tissue, while genes with a tau score closer to 0 are equally expressed across all tissues studied.

We downloaded 100 vertebrates conservation by PhastCons (phastCons100way) through the UCSC Genome Browser ^74^ (http://genome.ucsc.edu/). Subsequently, we used bigWigAverageOverBed ^74^ to extract the phastCons100way scores of CREs that constitute modular and non-modular networks.

#### Module hub CREs detection and associated CRE characteristic analysis

To identify CREs that act as connectors between regulatory modules, we proposed a hub CREs identification framework inspired by Guimerà et al. ^82^, which characterizes the topological role of each CRE using two complementary metrics: the participation coefficient *P* and the within-module degree *Z*.

The participation coefficient *P* quantifies how evenly a node distributes its connections across different modules and thus captures its inter-modular connectivity. For CRE *i, P*_*i*_ is defined as:

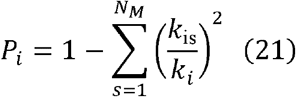

where *k*_is_ denotes the number of edges linking enhancer *i* to module *s, k*_*i*_ is the total degree of enhancer *i*, and *N*_*M*_ is the total number of modules in the network. Because the maximum attainable value of *P*_*i*_ depends on *N*_*M*_, we normalized it as:

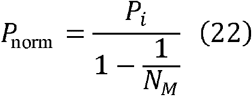

CREs with high *P*_norm_ frequently connect to multiple modules and are therefore candidates for inter-modular connectors.

To assess the importance of a CRE *i* within its own module, we computed the within-module degree *Z*_*i*_:

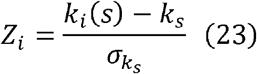

where *k*_*i*_ (*s*) is the number of connections of CRE *i* to other CREs in its assigned module *s, k*_*s*_ is the mean within-module degree of module *s*, and 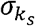 is the corresponding standard deviation. A high *Z*_*i*_ indicates that the CRE functions as a hub within its own module.

Therefore, CREs that exhibit both prominent inter-modular connectivity (with *P*_norm_ > 0.7) and intra-modular centrality (with *z*_*i*_ > 1) are defined as module hub CREs. This combined criterion identifies regulatory elements that not only bridge distinct regulatory modules but also occupy central positions within their own modules, consistent with a coordinating role in higher-order cis-regulatory organization.

We downloaded the transcription activity of K562 CREs quantified by MPRA experiment from a recent study conducted by Agarwal et al. ^23^. This study used lentivirus-based massively parallel reporter assays (lentiMPRAs) to test the regulatory activity of annotated cCREs among K562 and other two cell lines. The experiment verified K562 regulatory variant was downloaded from the recent study conducted by Siraj et al. ^38^. This study test the ability of 221,412 fine-mapped variants from complex and molecular traits to alter transcriptional output in K562 and other four cell lines using MPRA.

#### Comparison between inter- and intra-module interactions

The co-accessibility between CRE pairs was quantified by computing the product of their mean chromatin accessibility signals. The genome-wide Pol II-mediated ChIA-PET data of K562 is downloaded from the ENCODE portal (ENCSR880DSH) and used to evaluate the interaction strength between inter- and intra-module CRE pairs. To quantify the TF motif similarity between CRE pairs, we downloaded the cisTarget databases generated using the 2022 SCENIC+ motif collection ^53^, which is a matrix with CRE regions as rows, motifs as columns, and filled with values generated by scoring the DNA sequence of consensus peaks using Cluster-Buster. Subsequently, we employed this CRE-TF matrix to assess similarity in TF binding by calculating the cosine similarity between CRE pairs.

### Network dynamic analysis

#### Network dynamic score

We proposed a network dynamic score *Score*_dynamic_ to quantify the magnitude of network dynamics between healthy and disease states. *Score*_dynamic_ measures global changes in interaction strength. For each gene, ORIGAMI reconstructs weighted cis-regulatory interaction matrix 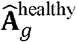 in healthy state and 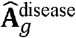 in disease state, where entries *A*_ij_ represent the predicted contact probability between regulatory elements.

We first computed the Frobenius norm of the difference between disease and healthy matrices:

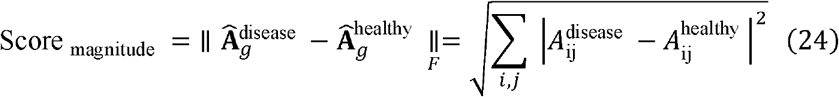

which captures the overall magnitude of network change regardless of direction.

To further distinguish whether a network predominantly gains or loses regulatory interactions, we decomposed the difference matrix

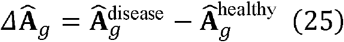

into two components. The positive difference matrix **P** retains only elements in 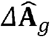 greater than zero, corresponding to interactions that are strengthened or newly gained in the disease state, with all other elements set to zero, and the negative difference matrix **N** retains only elements in 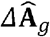 less than zero, corresponding to interactions that are weakened or lost in the disease state.

We then defined the direction of network dynamic score as:

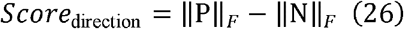

A positive *Score*_direction_ indicates that gained interaction strength outweighs lost interactions, reflecting an overall increase in regulatory complexity or activation in the disease state. In this case, the direction of *Score*_magnitude_ is set as positive; conversely, when *Score*_direction_ is less than zero, the direction of *Score* _magnitude_ is set as negative, yielding *Score*_dynamic_. The generated network dynamic scores were used to rank genes for the following gene set enrichment analysis using python package ‘gseapy’ ^41^.

#### Case study of network modularity change

We obtained the TF ChIP-seq for FOXA1 (ENCFF795BHZ) and ESR1 (ENCFF237WTX) on MCF-7 from the ENCODE portal. We downloaded 100 vertebrates conservation by phyloP (phyloP100way) through the UCSC Genome Browser ^74^ (http://genome.ucsc.edu/). We performed the survival analysis using Kaplan-Meier plotter ^83^, and the analysis results were downloaded and plotted using python package ‘lifelines’ ^84^.

#### Network dynamic cluster detection

To identify CRE clusters that exhibit similar covariation patterns of interaction changes with neighbors under different states, we drew inspiration from DiffCoEx ^85^, an algorithm designed to identify differentially co-expressed modules, proposed a network dynamic cluster detection method (details in Supplementary Methods). Intuitively, the method groups two genes together when their interactions to the same sets of genes change between the different conditions.

### In silico perturbation

#### MYC enhancer knockout

To validate the synergistic or additive effects of CRE interactions, we simulated enhancer knockout of the MYC cis-regulatory network at both the enhancer- and enhancer module-levels. First, we utilized ORIGAMI to construct the cis-regulatory network of MYC in K562, setting *τ* = 0.8, and only CREs that overlapped with the experimentally validated seven MYC enhancers (e1-e7) are retained for the following analysis. The louvain community detection algorithm divided the MYC cis-regulatory network into 3 modules: e1-e2, e3-e4, and e5-e7. Then, at the enhancer-level, we simulated single and paired enhancer knockouts by setting the DNase values of single or paired enhancers to zero. At the enhancer module-level, we clustered e1-e2 as c1, e3-e4 as c2, and e5-e7 as c3 and simulated single and paired enhancer module knockouts by setting the DNase values of single (e.g. e1-e2) or paired (e.g. e1-e2 & e5-e7) enhancer modules to zero. Last, ORIGAMI infer gene expression values and cis-regulatory network based on the perturbed input data. If the gene expression changes following paired perturbations exceed the sum of gene expression changes following two single perturbations (e.g. 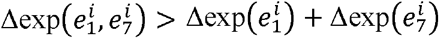 at enhancer-level or 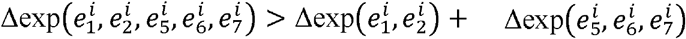 at enhancer module-level), we conclude that synergistic effects exist between inter-module CREs.

#### In silico eQTL variation and TF knockout

We employ ORIGAMI to predict gene expression changes induced by mutations in expression quantitative trait locus (eQTLs) and elucidate the mechanisms by which cis-regulatory variations influence gene expression. We downloaded the GTEx v8 CAVIAR fine-mapped eQTL analysis files from the GTEx portal ^52^ (https://gtexportal.org/home/downloads/) and selected SNP variants with CAVIAR posterior probability > 0.9 as fine-mapped eQTLs. We conducted the following analysis on fine-mapped eQTLs of ‘Breast_Mammary_Tissue’, as we collected sufficient epigenome profiles for mammary epithelial cell. The fine-mapped eQTLs with negative effect size (slope < 0) were classified as down-regulated eQTLs and those with positive effect size (slope > 0) were classified as up-regulated eQTLs. The fine-mapped eQTLs were introduced into the input genome sequences and used to predicted target gene expression change. Then, a gradient-based attribution analysis was conducted to interpret how noncoding mutations influence gene expression (details in Supplementary Methods).

For each gene in the K562 genome, we simulated TF knockout by changing input epigenome profiles of regulatory elements and predicted the gene expression after TF knockout (details in Supplementary Methods). We compared the consistency between the predicted expression change directions with the DEGs detected after TF knockout experiment. The gene expression change after TF perturbation experiments of K562 were derived from a previous work ^53^. To interpret how TF knockout alters gene regulation, we conducted gradient-based attribution analyses on both epigenomic and DNA features. For epigenomic features, we computed gradients of predicted gene expression with respect to input DNase signals and derived nucleotide-level contributions of DNase using the gradient × input method. The DNA attributions for reference and perturbed conditions were employed to identify regulatory motif whose contribution to gene expression was selectively diminished or enhanced upon TF knockout (details in Supplementary Methods).

## Code availability

The code and tutorials are available via GitHub at https://github.com/zjupgx/ORIGAMI.

## Acknowledgments

We thank the Information Technology Center and State Key Lab of CAD&CG, the Innovation Institute for Artificial Intelligence in Medicine, Zhejiang University for the support of computing resources. We also express our gratitude to group members for critical reading of the manuscript and providing constructive suggestions.

